# Simplification of ribosomes in bacteria with tiny genomes

**DOI:** 10.1101/755876

**Authors:** Daria D. Nikolaeva, Mikhail S. Gelfand, Sofya K. Garushyants

## Abstract

The ribosome is an essential cellular machine performing protein biosynthesis. Its structure and composition are highly conserved in all species. However, some bacteria have been reported to have an incomplete set of ribosomal proteins. We have analyzed ribosomal protein composition in 214 small bacterial genomes (< 1 Mb) and found that although the ribosome composition is fairly stable, some ribosomal proteins may be absent, especially in bacteria with dramatically reduced genomes. The protein composition of the large subunit is less conserved than that of the small subunit. We have identified the set of frequently lost ribosomal proteins and demonstrated that they tend to be situated on the ribosome surface and have fewer contacts to other ribosome components. Moreover, some proteins are lost in an evolutionary correlated manner. The reduction of rRNA is also common in bacteria with tiny genomes with deletions mostly occurring in free loops. Finally, the loss of the anti-Shine-Dalgarno sequence is associated with the genome reduction, the number of lost ribosomal proteins, and with the loss of bL9 and TF.

## Introduction

The ribosome is a universal biosynthesis machine present in all eukaryotes and prokaryotes. A bacterial ribosome is comprised of the small (30S) and large (50S) subunits which together form the 70S particle (Kurland 1972; Ramakrishnan 2002). The bacterial ribosome consists of multiple proteins (RP, r-proteins) and three rRNA molecules – 16S in the small subunit, 23S and 5S in the large subunit (Kurland 1972). The main catalytic functions of the ribosome, such as the peptide bond formation, mRNA decoding, and translocation of mRNA and tRNA after the peptide bond formation, are performed by ribosomal RNA (rRNA) (Green & Noller 1997; Nissen et al. 2000; Schmeing et al. 2003). Moreover, rRNA molecules also determine the ribosomal spatial organization providing sites for binding of the ribosomal proteins (Khaitovich et al. 1999). The ribosome of *Escherichia coli* contains 21 proteins in the 30S subunit (bS1–bS21) and 33 proteins in the 50S subunit (uL1–bL36) (Ban et al. 2014; Schuwirth et al. 2005; Kaczanowska & Rydén-Aulin 2007). The role of ribosomal proteins is to stabilize the ribosome and to regulate the ribosomal activity (Aseev and Boni 2011). While the key role in the protein biosynthesis is played by rRNA, the r-protein composition also tends to be conserved in most bacteria (Roberts et al. 2008). Moreover, 33 r-proteins are conserved among different domains of life (Roberts et al. 2008; Smith et al. 2008; Lecompte et al. 2002).

The protein composition of bacterial ribosomes has been studied intensively. The analysis of single-gene knockout mutants of *E. coli* has shown that these bacteria are able to grow without proteins bS6, uS15, bS20, bS21, uL1, bL9, uL11, and bL25, but in most cases the growth rate of these knockout mutants is reduced (Baba et al. 2006). In another study of an *E. coli* knockout collection, nine r-proteins (uL15, bL21, uL24, bL27, uL29, uL30, bL34, and uS17) have been found to be nonessential for survival in experimental conditions (Shoji et al. 2011). A similar study in *Bacillus subtilis* has identified 20 r-proteins non-essential to growth in experimental conditions (Akanuma et al. 2012). Based on this result, the smaller r-proteins were proposed to have been incorporated into the ribosome relatively recently in evolution, and hence to be less essential. A phylogenetic analysis of 995 completely sequenced bacterial genomes has shown that 44 r-proteins are strictly ubiquitous, proteins bS16, bL9, bL19, bL31, bL34, and bL36 are rarely missing, while bS21, S22, bThx, bL25, and uL30 are absent in a large fraction of bacteria (Yutin et al. 2012). A comparative analysis of the translation apparatus of *Mollicutes* has shown that five r-proteins (uS14, S22, bL7, bL25, and bL31) are missing in all studied genomes, uL30 is missing in almost all of them, and bS1 has been lost in seven independent events in different clades (Grosjean et al. 2014). Finally, an analysis of endosymbiotic bacteria with small genomes from a variety of phyla has demonstrated that they lack the largest fraction of r-proteins, as only 17 of 21 small-subunit and 16 of 32 large-subunit r-proteins are universally present in these bacteria (McCutcheon and Moran 2012). For example, *Candidatus* Tremblaya princeps, an endosymbiont of mealybugs (*Pseudococcidae* family), has only 139 genes suggesting little possibility to carry the complete translation apparatus (McCutcheon and Moran 2012). While the published lists of essential r-proteins vary to some extent, bL25, uL30, bL31, bS21, S22, and bThx have been consistently reported to be the least essential.

Less is known about whether there exist universal features common to these non-essential proteins, and how the r-protein loss is linked to the genome reduction. Here, we analyzed the r-protein composition in bacteria with small genomes from a variety of phyla. We have identified a set of frequently lost proteins and found two patterns of evolutionary correlative r-protein loss. A majority of frequently lost proteins have been shown to be located on the ribosome surface and to form less contacts with other ribosome components, compared to universally conserved r-proteins. We also show that the loss of the anti-Shine-Dalgarno sequence is associated with the genome reduction, the number of lost ribosomal proteins, and specifically with the loss of bL9 and TF.

## Results

### Some ribosomal proteins are missing in bacteria with small genomes

We re-annotated all r-proteins in 214 bacterial strains with small genomes (less than 1 Mbp) from 38 genera of the following phyla: *Proteobacteria, Bacteroidetes, Spirochaetes, Tenericutes*, and *Actinobacteria* (Supplementary Table S1, Materials and Methods). For that, we used 65 Pfam domains: 63 domains for the canonical set of ribosomal proteins, and two domains of the trigger-factor protein (TF), a ribosome-associated chaperone (Supplementary Table S2).

Our results (Supplementary Fig. S1) show that two proteins are absent in almost all considered strains, S22 (all strains) and bThx (present in *Candidatus* Walczuchella monophlebidarum only).

All except 11 r-proteins are lost in at least one strain from our dataset. At that, proteins bL9, uL24, bL25, uL29, uL30, bL32, bL34, bL36, bS21, and TF are lost frequently, as they are absent in at least 19 strains from a variety of phyla. This set later on is called frequently lost proteins. R-proteins of the small subunit are more likely to be retained than the large-subunit r-proteins (Supplementary Fig. S1). The only frequently lost small-subunit protein bS21 is absent in 54 strains from seven genera, while the most frequently lost protein from the large subunit, uL30, is absent in 138 genomes from 18 genera. All frequently lost proteins had been lost independently multiple times in bacteria (Fig. 1, Supplementary Fig. S2). The largest number of r-protein losses was observed in *Candidatus* Tremblaya princeps, *Candidatus* Hodgkinia cicadocola, and *Candidatus* Carsonella rudii, bacteria with the shortest genomes in our dataset.

**Figure 1.**
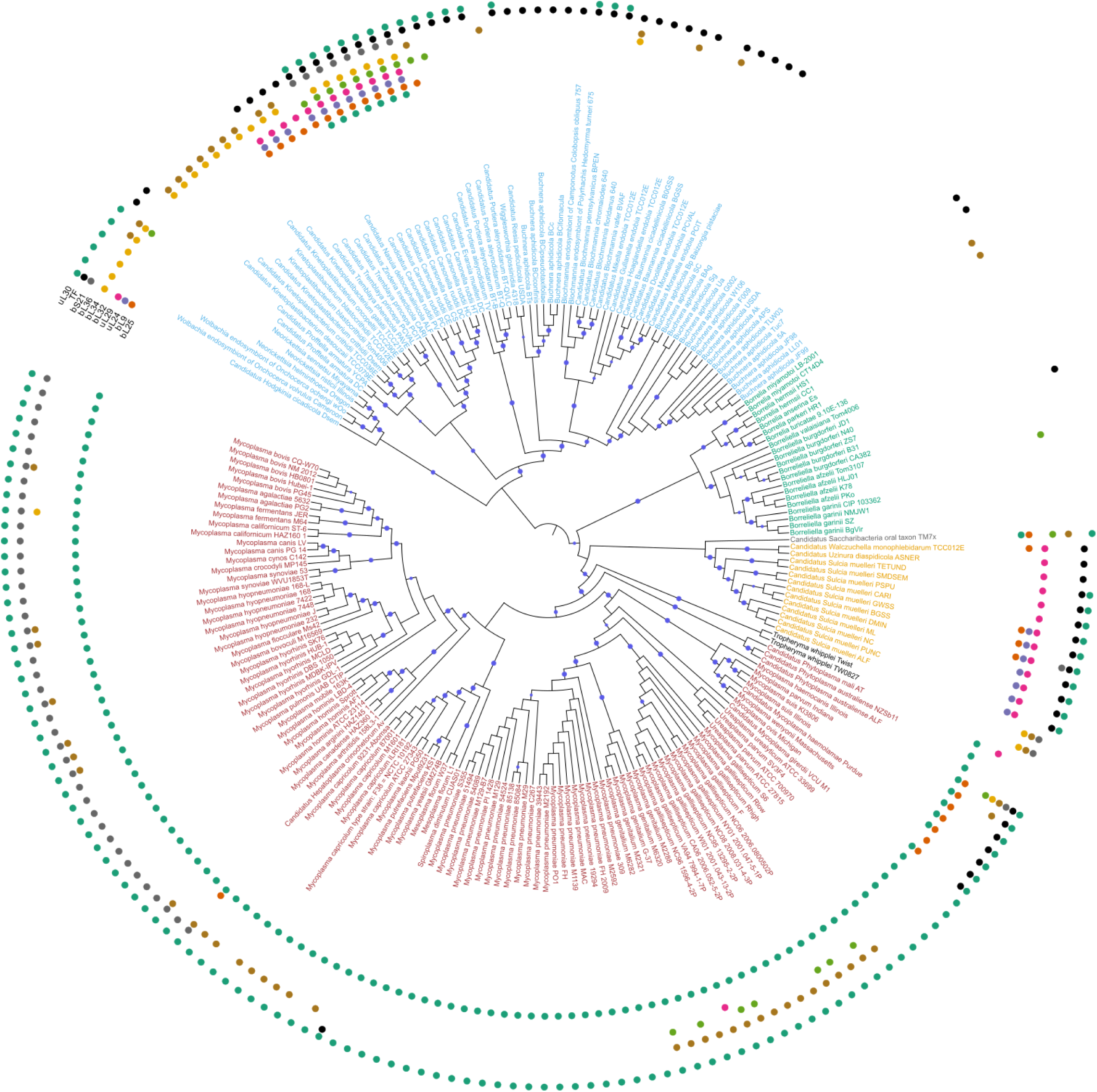
Maximum likelihood phylogenetic tree of analyzed bacterial species, constructed by concatenated alignment of conserved r-proteins. The tree was constructed by PhyML with 100 bootstrap replicates. In this representation branch lengths are ignored (the tree with branch lengths is provided as Supplementary Fig. S2). Bootstrap values in the range 0.9–1 are shown by blue circles. The absence of one of ten frequently lost proteins (bL9, bL21, L24, bL25, uL29, bL32, bL34, bL36, bS21, TF) in a strain is marked by a colored circle. Leaves are colored by the phyla: *Actinobacteria* – black, *Bacteroidetes* – yellow, *Proteobacteria* – blue, *Spirohaetes* – green, *Tenericutes* – red, unclassified bacteria – grey.

### R-protein loss depends on the level of genome reduction

The comparison of the r-protein composition and the genome size revealed a correlation between the genome size and the number of retained ribosomal proteins (*r*^2^ = 0.6, *p* < 2.2×10^−16^), but the slope is dramatically different for the tiny genomes (Supplementary Fig. S3a). Indeed, for genomes shorter than 350 Kb there is an even stronger correlation between the genome size and the number of r-proteins (*r*^2^ = 0.7, *p* = 5.7×10^−6^), while the remaining genomes in the dataset show no correlation at all (*p* = 0.76). This pattern holds both for the complete ribosome and the individual subunits (Supplementary Fig. S3bc), the slope being steeper for the large subunit.

### Patterns of ribosomal protein loss

Ten most frequently lost proteins have been lost independently in different phyla, showing parallel events (Fig. 1, Supplementary Fig. S2). A general tendency is that frequently lost proteins are usually situated at the ribosome surface (Fig. 2). To analyze possible dependencies between frequent losses, we estimated the correlation between the vectors of the protein presence/absence in all strains by the Pagel correlation method (Pagel 1994) which allows one to control for the phylogenetic structure of the dataset (Fig. 2a). After the Bonferroni correction two clusters were observed, uL29+TF and bL32+bL34+bL36.

**Figure 2.**
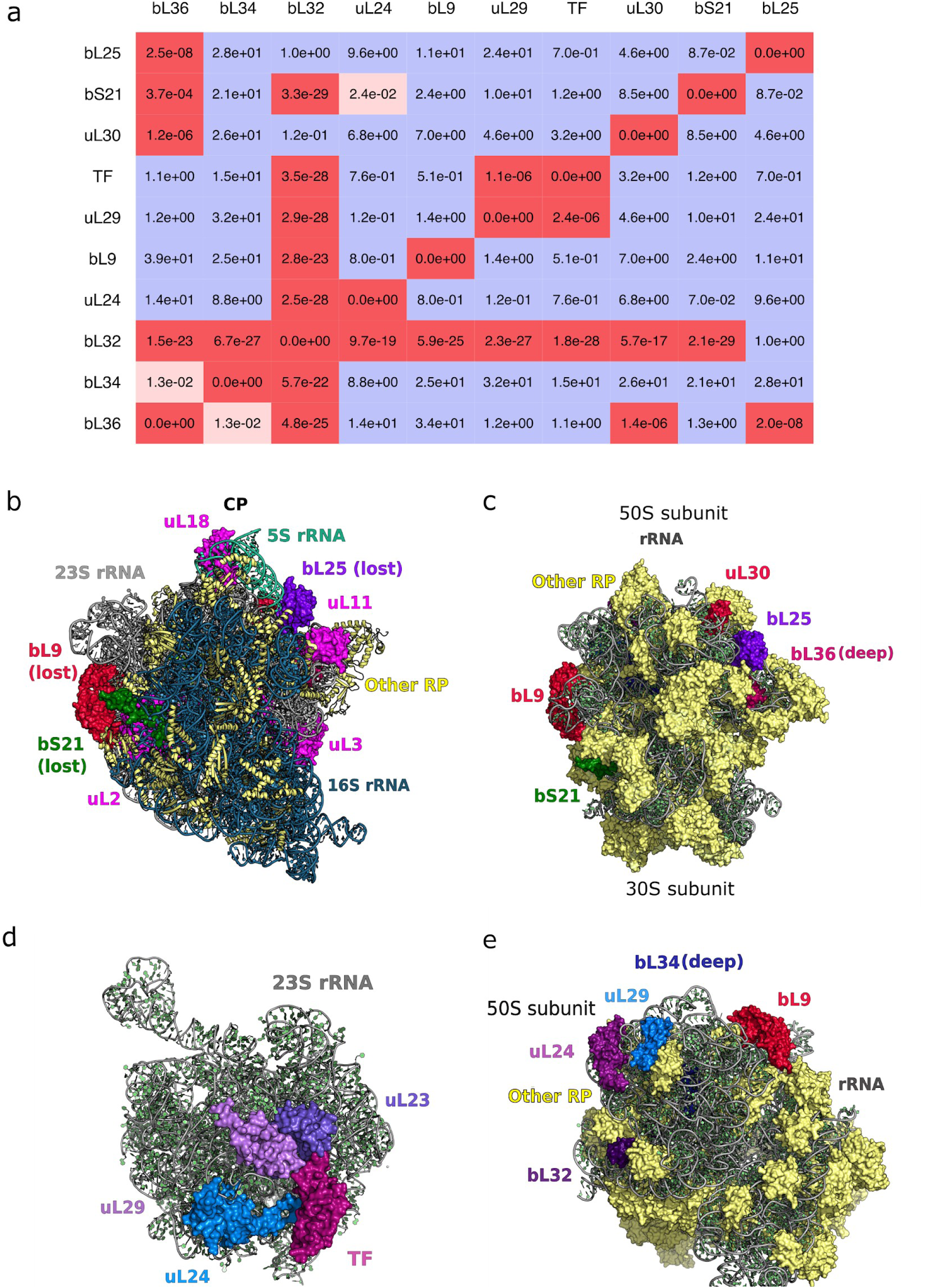
Patterns of r-protein loss. (a) Pagel’s test of correlated evolution between vectors of ribosomal protein presence/absence in bacterial strains with the Bonferroni correction. Insignificant p-values are colored blue, the significant ones are colored red (the more significant, the more intense). Only ten frequently lost proteins are considered. (b) The crown view of the 70S ribosome (the typical crown view position is marked using several ribosomal proteins (magenta), the position of the central protuberance and 5S rRNA (light green) and the position of 16S rRNA (dark blue)) and various perspectives (c,e) of the ribosome showing positions of frequently lost proteins. PDB ID: 5H5U (*Escherichia coli*) (Ma et al. 2017). (d) 23S rRNA and r-proteins which are connected with trigger factor. PDB ID: 2D3O (*Deinococcus radiodurans*) (Schlünzen et al. 2005). Protein labels are of the same color as the respective proteins in the structure. In (c), (e) only frequently lost ribosomal proteins are labeled.

One of the observed loss clusters is consistent with the r-proteins location in the ribosome. The correlated loss of proteins uL29 and TF may be explained by the TF position in the ribosomal structure, as the latter is located at the ribosome surface where it is surrounded by r-proteins uL23, uL24, and uL29 (Fig. 2d) directly interacting with uL23 and uL29 (Ferbitz et al. 2004). The second pattern may not be explained by the structural proximity as bL36 is situated far from bL32 and bL34. These r-proteins share a common feature of being deeply immersed within a structure, with bL32 having a surface region, while bL34 and bL36 being immersed completely (Fig. 2bce).

In the “onion model” of the ribosome structure and evolution (Hsiao et al. 2009; Petrov et al. 2015), the peptidyl transferase center (PTC) is the core and the most ancient part of the ribosome. According to this model, the farther is an r-protein from the PTC, the later it has been added during ribosome evolution. Indeed, most of the frequently lost proteins are situated in the outer layers of the “onion”, except for the r-proteins bL34, bL36, bL25, uL30. However, there is no significant difference between conserved and frequently lost r-proteins in the distribution of distances from the PTC.

### Frequently lost r-proteins have less contacts than conserved proteins

To study what features differ between frequently lost and conserved r-proteins, we fitted a logistic regression model to our data, regressing the protein status (either conserved or frequently lost) on the evolutionary rate, protein length, and the number of contacts with all adjacent proteins and rRNA. Only the number of contacts was a significant determinant of the protein status (*p* = 0.022).

We further analyzed the differences in the evolutionary rate (Fig. 3a) and the number of contacts (Fig. 3b) between conserved and frequently lost proteins (see Supplementary Table S3). Indeed, frequently lost proteins form significantly less contacts than conserved proteins (means 231 and 478, respectively, *p* = 1.4×10^−4^), as the former are mainly situated on the ribosome surface. These findings are supported by measurements of the contacting surface between each of the studied r-proteins and the rest of the ribosome (the Wilcoxon test *p* = 2.0×10^−24^ for PDB structure 5H5U that does not contain TF, *p* = 8.2×10^−5^ when the value for TF from PDB structure 2D3O is added to data for PDB structure 5H5U). While several PDB structures of the ribosome are available, none of them is complete (i.e. includes the entire ribosome and TF) and for none all parts of the ribosome are resolved. Because of that we considered several PDB structures, however, the results do not depend on the selected structure (Supplementary Fig. S4). However, if the contacting surface values are normalized on the protein’s surface, the difference between frequently lost and conserved r-proteins becomes insignificant, and hence the absolute interacting surface is important, and not a relative one.

**Figure 3.**
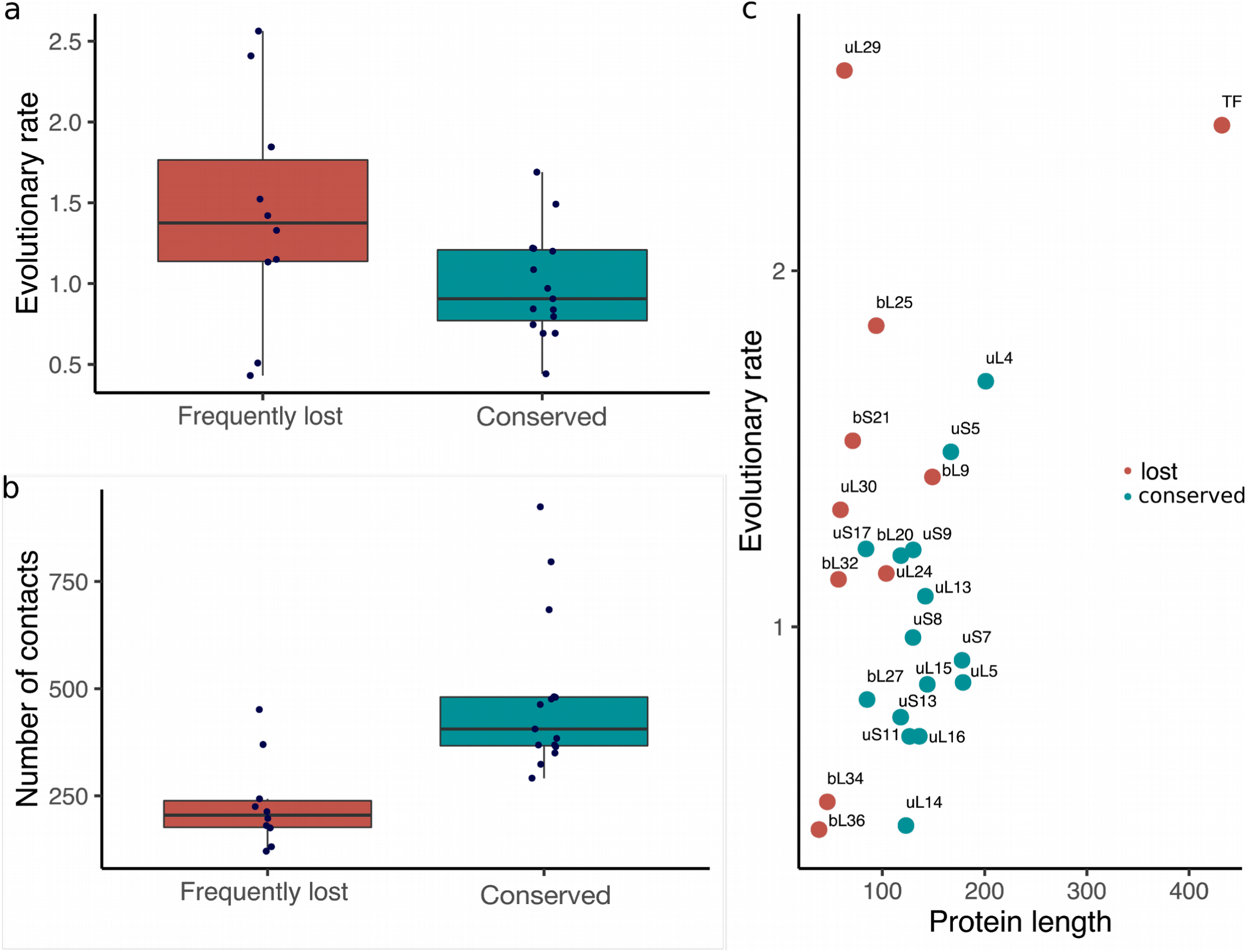
Determinants of the r-protein loss. The differences in the evolutionary rate (a) and the number of contacts (b) between the conserved (blue) and frequently lost (red) ribosomal proteins. (c) The dependency between the evolutionary rate and protein length (number of amino acid residues) for both conserved and frequently lost ribosomal proteins.

Surprisingly, there was no significant difference in the evolutionary rate between conserved and frequently lost proteins. Moreover, the evolutionary rate was slightly correlated with the protein length (*r*^2^ = 0.39, *p* = 0.055), with short proteins evolving slower (Fig. 3c).

### Deletions in rRNAs and the loss of r-proteins

To study possible correlations between the loss of r-proteins and their rRNA binding sites, we analyzed 16S and 23S rRNA multiple alignments and identified nine deletion blocks (four in 23S and five in 16S) that had occurred in more than two genera and were longer than five nucleotides in at least one species. All such deletions affect rRNA free loops not involved in the r-protein binding (Supplementary Fig. S5).

### Anti-SD loss is frequent among strains from different taxa

The anti-Shine-Dalgarno sequence in 16S rRNA is frequently lost in diverse phyla. Its presence/absence pattern is not strongly linked to the taxonomy, as many phyla have strains both with and without the anti-SD sequence in 16S rRNA (Supplementary Fig. S6). The loss is significantly associated with the genome reduction (one-sided Wilcoxon test *p* = 0.034) and with the number of lost r-proteins (one-sided Wilcoxon test *p* = 1.04×10^−5^). We have observed a strong correlation of the anti-SD loss with loss of TF and some correlation between the losses of anti-SD and bL9, but with lower confidence (Pagel’s test with the Bonferroni correction *p* = 7.55×10^−5^ and *p* < 0.05, respectively; see Supplementary Table S4).

## Discussion

The bacterial genome size is evolutionarily labile, and radical genome reduction has occurred many times independently throughout the bacterial domain. Such multiple independent genome reduction events allows for a systematic study of the patterns of gene loss associated with them.

While individual r-proteins can be lost in bacteria with genomes of any size, for the tiniest of bacterial genomes, further reduction in the genome size is associated with the loss of generally conserved ribosomal proteins. The fact that ribosomal proteins are among the last to leave a shrinking genome illustrates that they tend to be less dispensable than most proteins. Two r-proteins absent in the largest number of small-genome bacteria are S22 and bThx. Previous studies indicate that these proteins are largely non-essential. Indeed, the gene encoding S22 is mostly expressed at the stationary growth phase, and its deletion does not affect the viability of *E. coli* mutants (Izutsu et al. 2001). bThx is found only in thermophilic bacteria and has been shown to stabilize the organization of RNA elements at the top of the 30S subunit head in *Thermus thermophilus* (Choli et al. 1993; Wimberly et al. 2000), which implies that this protein is only essential for survival at high temperatures. However, the low prevalence of these two proteins may indicate not the ease of their loss, but rather their late arrival. S22 is present only in some enterobacteria (Izutsu et al. 2001), and bThx homologs are only present in thermophilic bacteria from genus *Thermus* (Choli et al. 1993; Wimberly et al. 2000). Therefore, both S22 and bThx seem to have been acquired only by some bacterial groups long after the last common ancestor of bacteria.

Our list of frequently lost proteins is largely consistent with previous observations (Baba et al. 2006; Akanuma et al. 2012; Yutin et al. 2012; Grosjean et al. 2014; Shoji et al. 2011). For example, among six non-ubiquitous r-proteins reported by Yutin et al. 2012, three (bL25, uL30, bS21) are also among the r-proteins identified as frequently lost in our dataset, in addition to two proteins (S22 and bThx) that only occur in some bacterial lineages (see above). However, our focus on bacteria with tiny genomes allowed us to study less dispensable r-proteins. Proteins bL34 and bL36 identified previously as present in almost all genomes (Yutin et al. 2012) are frequently lost in our dataset. While these proteins are generally considered essential, they can be dispensable under certain circumstances. The absence of bL34 affects cell growth, but the cell function can be restored by increasing the magnesium ion flow (Shoji et al. 2011; Akanuma et al. 2014). The absence of bL36 is only essential for cell growth at high temperatures (Ikegami et al. 2005). Moreover, bL36 is missing in *Bacteroidetes* (Yutin et al. 2012). R-proteins uL24 and uL29 identified here as frequently lost had been lost in at least two independent events, and only in bacteria with drastically reduced genomes. The knockouts of both these proteins are viable (Shoji et al. 2011). Interestingly, r-proteins bL31 and uS14 have been identified as lost in all *Mollicutes* (Grosjean et al. 2014), but we have observed that only few strains of *Mollicutes* lack these proteins. This difference is likely caused by a more sensitive annotation procedure used here.

Extreme genome reduction is common not only in endosymbiotic bacteria but also in cell organelles, mitochondria and plastids. The r-protein composition of these organelles has been extensively studied (Yamaguchi and Subramanian 2000; Maier et al. 2013; Petrov et al. 2018). The plastid r-proteins core encoded by all plastids is comprised of only 15 r-proteins of bacterial origin (Maier et al. 2013). This set contains only one of the frequently lost r-proteins reported here, bL36, while other frequently lost r-proteins are frequently absent in plastids (Maier et al. 2013). The protein composition of mitochondria is highly variable across species and is characterized not only by gene loss but also by acquisition of proteins of various origins (Petrov et al. 2018). Among eight large-subunit proteins in our list of frequent losses, six r-proteins (bL9, bL25, uL30, bL32, bL34, bL36) are absent in mitochondria in at least one studied species, but some proteins of bacterial origin are not simply lost but are replaced by non-homologous mitochondria-specific ones. Moreover, while we have observed frequent loss of uL24, in mitochondria this r-protein is not only universally conserved but additionally stabilized by mitochondria-specific mL45 (Petrov et al. 2018). In any case, while the patterns of loss seem superficially similar, plastids, mitochondria, and bacteria with reduced genomes cannot be compared directly, since a large fraction of the organellar genome has been transferred to the nuclear genome. This leads to a different selection regime and the absence of the selective pressure on the proteome size. For example, spinach chloroplast contains 59 r-proteins, of which 53 are orthologous to *E. coli* ones. Of those, some r-protein genes are still retained in the plastid genome, this groups being comprised of about half of small subunit r-proteins (12 out of 25) and eight out of 33 large subunit r-proteins (Yamaguchi and Subramanian 2000; Yamaguchi et al. 2000). Among frequently lost proteins reported here, r-proteins bL25 and bL30 are missing in spinach, two more are encoded in the spinach chloroplast genome (bL32 and bL36), while bS21, bL9, uL24, uL29, and bL34 are encoded by the nuclear genome.

What are the factors that affect the propensity of a protein to be lost? Previously, proteins frequently lost in evolution have been shown to evolve rapidly, have fewer interactions with other proteins, and lower expression levels (Krylov et al. 2003). Although there are some data on ribosomal heterogeneity in bacteria (Byrgazov et al. 2013), the complete ribosome requires all r-proteins, and r-proteins are organized in operons with a variety of regulatory feedback loops providing tight co-regulation (Lemke et al. 2011), that allowed us not to consider the expression level. Taking into account other factors such as the substitution rate, the number of contacts, and the protein length, we observe that only the number of contacts is a significant determinant of the r-protein loss. The number of contacts is associated with exposure of an r-protein on the ribosome surface. Indeed, r-proteins that are frequently lost tend to be located on the ribosome surface, the exceptions being bL34 and bL36 (Fig. 2ce).

The order in which r-proteins are assembled in the ribosome complex (Chen & Williamson 2013) provides an indirect evidence of the order in which r-proteins have been incorporated in the ribosome in the course of evolution. Thus, r-proteins that appear only in the late ribosome intermediates should be relatively young. Seven out of ten frequently lost proteins r-proteins – bS21, bL9, bL25, uL29, bL32, bL34 and bL36 – are included only in the late assembly intermediates. Thus, it can be speculated that frequently lost r-proteins have been incorporated in the ribosome late in evolution. This hypothesis is also supported by “the onion model” (Hsiao et al. 2009; Petrov et al. 2015). Most frequently lost proteins in our dataset are located at the farthest layers from the PTC and thus, they should have appeared the latest in the evolution. Surprisingly, r-proteins bL25, bL34, and bL36 are closer to the PTC then other frequently lost r-proteins, however they are included in the late assembly intermediates (Chen & Williamson 2013). These proteins possibly appeared late in evolution, but then migrated deeper in the ribosome structure. Another interesting question is whether r-proteins are lost at random or there are certain patterns of ribosome simplification, where the loss of one protein leads to the loss of others. Our dataset contained many genomes with incomplete sets of r-proteins, that allowed us to study the patterns of common protein loss.

One such pattern is the loss of proteins around the ribosome exit tunnel (TF, and uL29). The trigger-factor (TF) is a chaperone associated with the ribosome exit channel (Hoffman et al. 2010), and is the most frequently lost protein in the pattern. TF is directly interacting with uL23 and uL29 (Ferbitz et al. 2004). We show that the loss of uL29 is correlated with the loss of TF, while the loss of uL23 is not associated with TF. The loss of this functional block is surprising, as even highly reduced genomes tend to retain chaperones (McCutcheon and Moran 2012).

One more pattern is the correlated loss of bl32, bL34, and bL36. Two proteins in this pattern – bL32 and bL34 – are positioned near r-proteins forming the exit tunnel, uL22, uL23, uL24, and uL29 (Nikolay et. al 2015), whereas bL36 is far from bL32 and bL34, so this pattern may not be explained just by the spatial proximity. The only common structural feature of these three proteins is that they all are buried deeply in the structure. At that, little is known about the functional role of these r-proteins. bL32 together with bL17 determine the LSU assembly path and without these two r-proteins the subunit core becomes unstable (Davis et. al 2016; Nikolay et. al 2018). bL36 has been reported to be highly conserved in Bacteria, but lacking in Archaea and Eukarya (Maeder and Draper 2005). bL36-deficient ribosomes exhibited disruptions of the rRNA tertiary structure, suggesting that bL36 is important for the 23S rRNA organization and stabilization (Maeder and Draper 2005). Hence, this entire group may be associated with the 23S rRNA stabilization.

Among other frequently lost proteins, uL30 is highly conserved in Archaea and Eukarya (Lecompte et al. 2002), where it is thought to be essential for the selenocysteine recognition (Chavatte et al. 2005). The role of this protein in bacteria is not clear. bL25 has been proposed to be essential for interaction with r-protein uL16, the latter being necessary for the ribosome stability (Anikaev et al. 2016). Although the loss of bL25 is tolerated in *E. coli*, mutant bacteria have a reduced growth rate (Baba et al. 2006). bS21 is required for the recognition of native templates, and its function resembles the function of bS1 (Van Duin and Wijnands 1981). R-protein bL9 reduces translation frameshifting (Dunkle et al. 2010). uL24 is an LSU assembly initiator protein, together with uL3 (Nikolay et. al 2015). TF and uL29 discussed in detail above are associated with protein folding. Thus, the loss of these proteins should reduce the ribosome fidelity and overall efficacy of protein translation.

Surprisingly, all deletions in 23S and 16S rRNAs happened in free loops meaning that the loss of ribosomal proteins was not accompanied by the reduction of their respective binding sites on the ribosomal RNAs. However, the shortening of rRNA is a tendency shared also by mitochondrial rRNA — the SSU mt-rRNA is 40% shorter than bacterial 16S rRNA, while LSU mt-rRNA represents a half of bacterial 23S rRNA (Anderson et al. 1981). Interestingly, deletions in 23S rRNA that have been found in our dataset are located in helices h10, h63, h79, and h98, which have been recently reported to be lost frequently in mt-rRNA of various species (Petrov et al. 2018). Degeneration and losses of a particular element of 16S rRNA, the anti-Shine-Dalgarno sequence, have been reported in various species including symbionts with small genomes (Lim et al. 2012). The anti-SD loss is polyphyletic, as the sequence may be present or absent in representatives of all considered phyla, consistent with previous observations (Amin et al. 2018). While the loss of the anti-SD sequence is a common event among intracellular symbionts, some symbionts with reduced genomes, such as *B. aphidicola*, retain anti-SD (Lim et al. 2012). Nevertheless, we have observed that the loss of anti-SD is significantly associated with the genome reduction and the loss of ribosomal proteins. Moreover, we see an association between the loss of anti-SD and the absence of frequently lost ribosomal proteins bL9 and TF. Surprisingly, none of them are situated near the 3’ end of 16S rRNA, where the anti-SD sequence is located. Thus, the association between the absence of these proteins and the anti-SD sequence may not be explained by direct interactions, and changes in structural dynamics should be taken into account.

In conclusion, proteins are not lost at random, and there are at least two distinct patterns of loss of spatially adjacent proteins. Overall, frequently lost proteins have less contacts, are located on the ribosome surface, and have been incorporated in the ribosome late in evolution. That is consistent with loss not affecting essential ribosome functions, especially in symbiotic bacteria that live in stable host environments.

## Materials and Methods

### Dataset of small bacterial genomes

The list of all bacterial species with complete genomes not exceeding 1 Mbp was compiled from the IMG/M database (Chen et al. 2017). The genomic data for all strains of these species (214 genomes in total, Supplementary Table S1) were downloaded from the NCBI FTP; files with protein sequences in February 2017 (ftp://ftp.ncbi.nlm.nih.gov/genomes/refseq/bacteria/), files with RNA sequences in September 2017 (ftp://ftp.ncbi.nlm.nih.gov/genomes/genbank/bacteria/).

The genomes of some strains in the selected species exceeded 1 Mb, but were still retained in the dataset to check whether the gene loss tendency is consistent throughout the species.

### Annotation of protein domains

Ribosomal proteins and trigger factor (TF) were re-annotated in the downloaded genomes (the complete list of proteins is given in Supplementary Table S2) using HMM matrices from the Pfam-A database (Finn et al. 2016). The HMM-profiles for the selected domains were scanned against each genome with the HMMER software (Mistry et al. 2013). A protein was considered to be present in a given genome if the respective domain had a hit in the HMMER search with E-value < 0.001; only the best such hit was retained for further analysis. If the HMMER domain bias had the same order of magnitude as the similarity score, additional filtering was performed as described below.

For each domain we created a multiple alignment of proteins with confident HMMer predictions (available at https://github.com/darianick/ribo-simpler). Protein alignments were built using the GUIDANCE2 Server (Penn et al. 2010) with the MAFFT program for building multiple sequence alignments. A candidate protein with a high domain bias value was added to the domain alignment and considered as present if it fitted the multiple sequence alignment keeping the GUIDANCE alignment confidence score above the recommended threshold (0.8). If the confidence score was below the threshold, the protein was removed from the dataset.

Two-domain proteins (uL2, uL5, bL7/L12, bL9, uL11, uS4, uS5, TF) were considered to be present if both domains were found.

### Identification of independent r-protein losses

To map the phyletic patterns of r-protein losses, we constructed the maximum likelihood tree of concatenated multiple alignments of all conserved r-proteins (see below) using PhyML v. 3.1 (Guindon et al. 2010) with the default parameters and 100 bootstrap replicates (available at https://github.com/darianick/ribo-simpler). Prior to the tree construction all columns containing gaps were removed from the concatenated alignment. This tree was used to calculate the number of independent losses of each protein.

### Estimation of the evolutionary rate

The evolutionary rate of a protein was defined as the average branch length in the respective phylogenetic tree. The maximum likelihood phylogenetic tree was built for each ribosomal protein using PhyML with 100 bootstrap replicates (LG+G model was utilized with tree topology and branch lengths optimised (“-o tl” parameters) and with gamma parameter identified for each protein alignment using ProtTest (version 3.4.2) (Darriba et al. 2011)). The evolutionary rate for a protein was compared to the average rates for the set of conserved proteins (uL5, uL14, uL15, bL20, uS5, uS7, uS8, uS9, uS11, uS13, and uS17). For each studied protein, the corresponding rates for every conserved protein from the set were calculated in the subtree with the same set of species as in the tree of the studied protein. To compare the results for proteins with different phyletic profiles, we calculated the relative evolutionary score, i.e. the protein’s evolutionary rate divided by the corresponding average rate for the set of conserved r-proteins. For two-domain proteins (see Supplementary Table S5) the same gamma parameters and, subsequently, the same evolutionary rates were obtained, with the exception of ribosomal protein bL9 - for this protein we utilize an average evolutionary rate between the two evolutionary rates for domains.

### Detection of patterns of ribosomal protein loss

The p-values for pairwise correlations between losses of r-proteins were estimated using Pagel’s test for correlated evolution (Pagel 1994) with R-package phytools v0.6-99 (Revell 2012) based on vectors of ribosomal protein presence/absence in all studied bacterial strains and the phylogenetic tree of conserved r-proteins (see above). The resulting p-values after the Bonferroni correction were organized in a heatmap with significant hits grouped together and representing patterns of evolutionary correlated r-protein loss. The obtained results are not always stable (e. g. bS21 vs. uL24, uL30 vs. bL32, bS21 vs. bL36 on Fig. 2a) due to the presence of long branches on the tree.

### Calculation of the number of contacts

The number of contacts was estimated by measuring pairwise distances between atoms in ribosome PDB structures (PDB ID: 5H5U and 2D3O) (Ma et al. 2016; Schlünzen et al. 2005). Atoms were defined as contacting if the distance between them was at most 5Å. All protein self-contacts were disregarded. When multiple atoms of a studied protein were connected with the same atom in the ribosome structure, only one such contact was retained. The pairwise distances were calculated with the PyMOL Molecular Graphics System, Version 1.8 Schrödinger, LLC and script pairwise_dist.py. For each protein the total number of contacts was calculated. The contacting surface between each studied r-protein and the remaining ribosome was calculated using the “get_area” function in PyMOL by the following formulae:

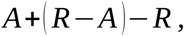

where A is a r-protein surface, R is a surface of a complete ribosome, R - A is a surface of the complete ribosome lacking r-protein A.

The distance between r-proteins and PTC was measured using “distance” function in PyMOL with a parameter “mode = 4” to obtain the distance between centroids of r-proteins and PTC. PTC was set as residues 2063-2663 of 23S rRNA.

### rRNA analysis

16S and 23S rRNA alignments (available at https://github.com/darianick/ribo-simpler) were built using SINA Alignment Service (Pruesse et al. 2012), and then all common gaps were removed. We considered rRNA deletions if they occurred in more than two genera and were longer than five nucleotides in at least one species. Two deletion blocks where deletions were present at the same set of strains and separated by not more than five nucleotides in the reference sequence were considered as a single deletion block. As a reference we selected 23S and 16S rRNA sequences from PDB ID: 5H5U.

A strain was defined as having the anti-Shine-Dalgarno sequence if there was an anti-SD motif CCUCCU at the 3’-end of the strain’s 16S rRNA.

### Statistical analysis and data visualization

All statistical analyses were performed using R. We used the logistic regression (function glm, family=binomial()) to find determinants of r-proteins loss. The correlation test was performed in R with the cor.test() function. The Wilcoxon(-Mann-Whitney) unpaired rank sum test was performed using “wilcox.exact()” function from “exactRankTests” R package. The dependency between the total number of r-proteins and the genome length (Supplementary Fig. S3) was built in R using the ggplot2 package. Curve smoothing was performed using the “loess” method. Phylogenetic trees were visualized with the iTOL server (Letunic and Bork 2011).

## Acknowledgements

The project was initiated with Sofya Sherstneva and Anna Ergemlidze at the Summer School of Molecular and Theoretical Biology for high school students, supported by the Zimin Foundation. We thank Arthur Zalevsky (FBB MSU) and Georgii Bazykin (IITP, Skoltech) for useful discussions and suggestions. This work was supported by the Russian Science Foundation (grant 18-14-00358).

