## Supplementary figures and tables for "Simplification of ribosomes in bacteria with tiny genomes"

**Figure S1. Heatmap showing the presence of ribosomal proteins in strains of bacteria with small genomes.** Ribosomal proteins are in rows; genomes are in columns, clustered by phyla. Absent ribosomal proteins are shown by dark blue. Frequently lost proteins are labeled in red.

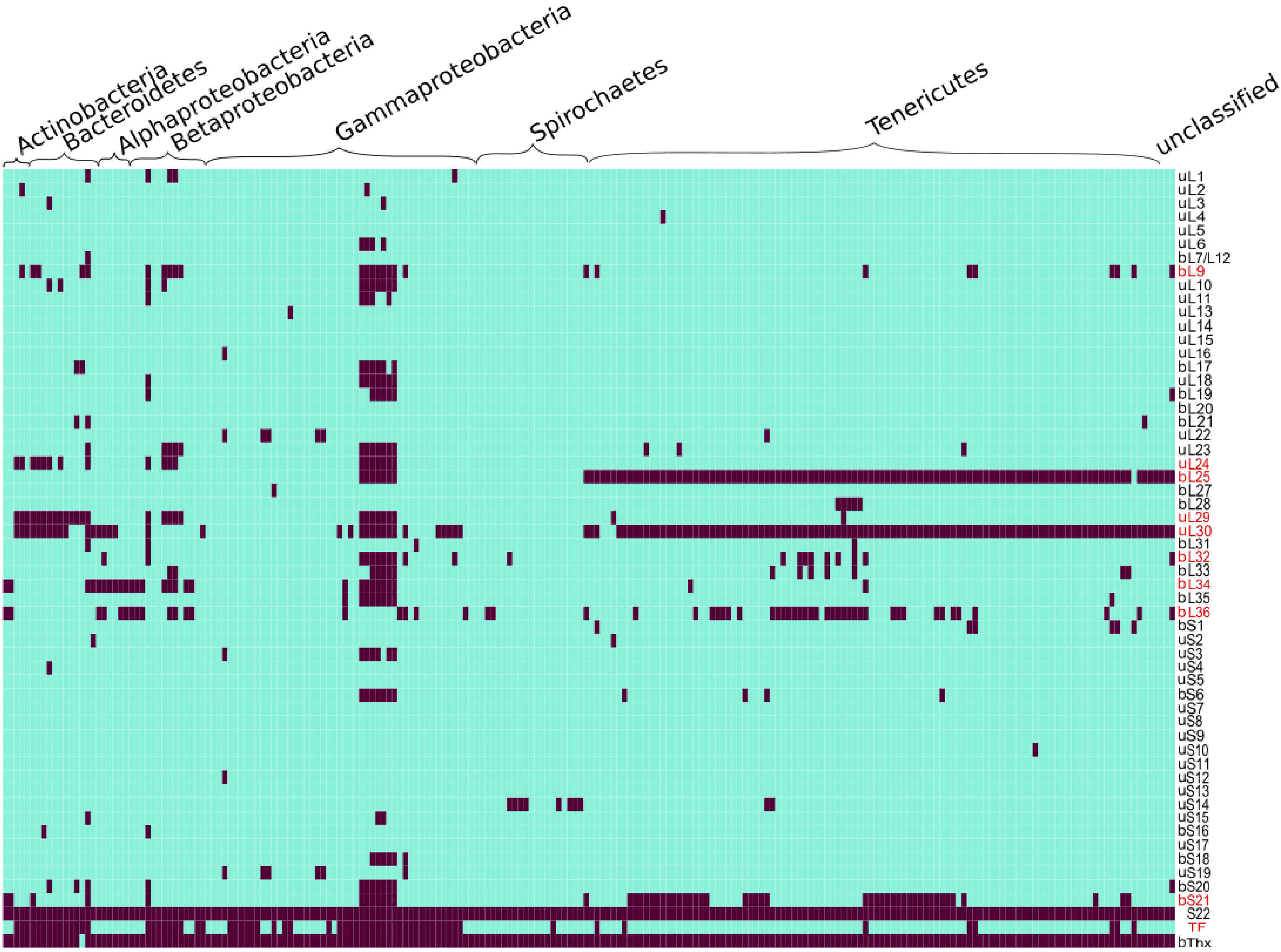

**Figure S2. Maximum likelihood phylogenetic tree of analyzed bacterial species, constructed by concatenated alignment of conserved r-proteins.** The same tree is provided in Fig. 1, but here the branch lengths are provided. Bootstrap values in the range 0.9–1 are shown by blue circles. The absence of one of ten frequently lost proteins (bL9, uL24, bL25, uL29, uL30, bL32, bL34, bL36, bS21, TF) in a strain is marked by a colored circle. Leaves are colored by phyla: *Actinobacteria* – black, *Bacteroidetes* – yellow, *Proteobacteria* – blue, *Spirohaetes* – green, *Tenericutes* – red, unclassified bacteria – grey.

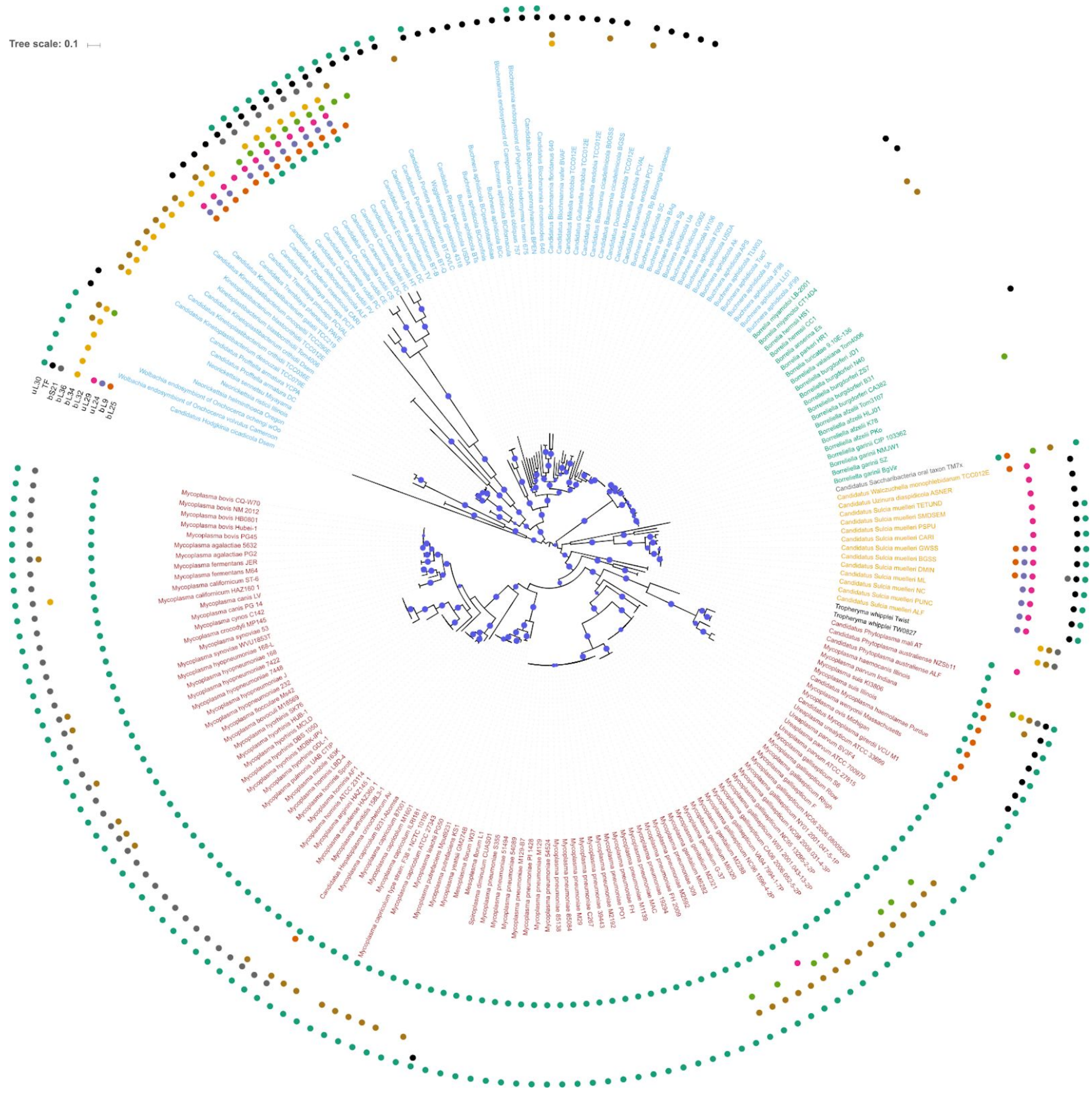

**Figure S3. Dependencies of the number of ribosomal proteins (RP) on the genome length** (in bp): (a) in the ribosome, (b) in the 50S subunit, and (c) in the 30S subunit. Each dot represents a bacterial genome, the dots are colored by phyla listed in the legend. The trend line is generated by the loess regression.

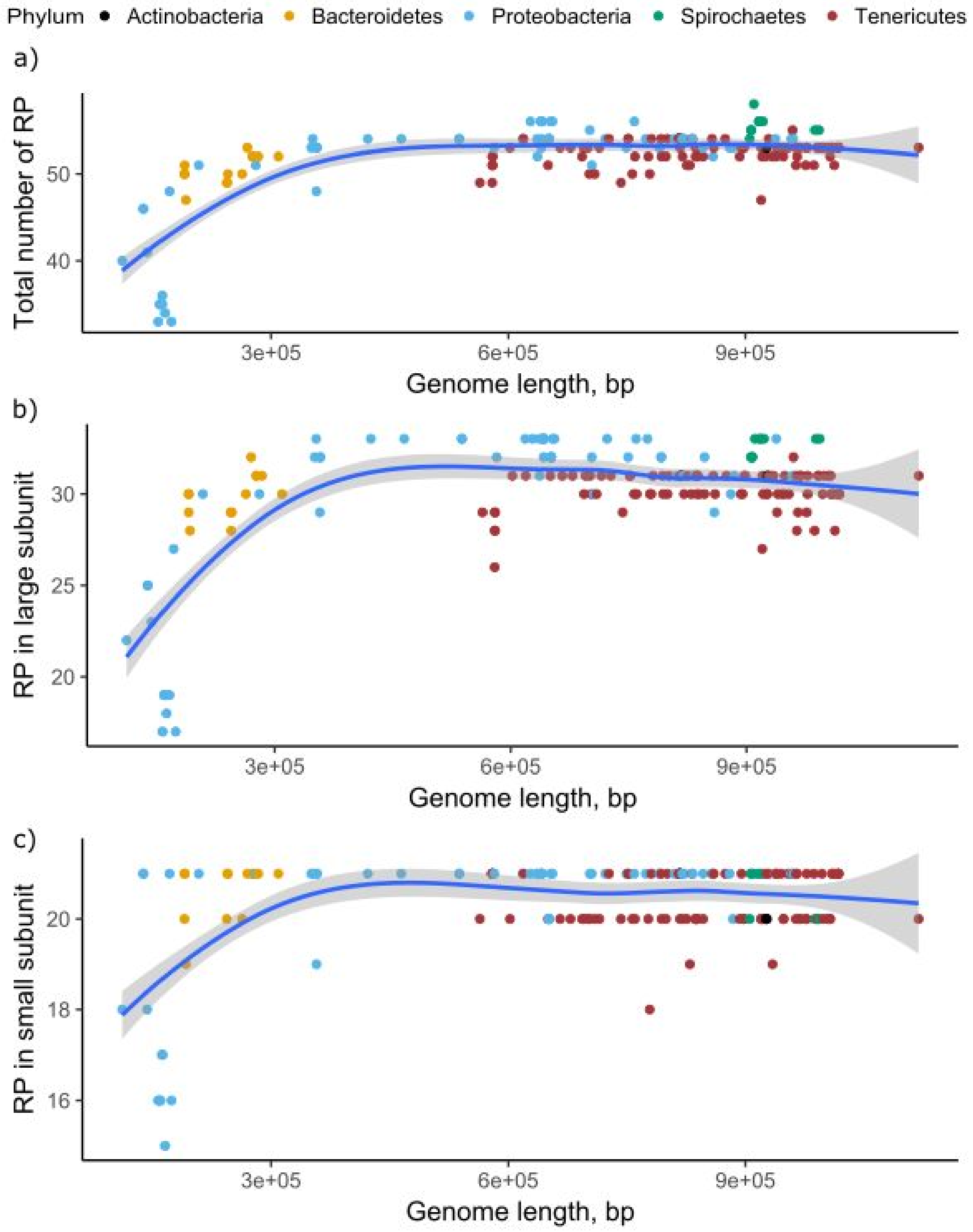

**Figure S4. Contact surface comparison between frequently lost and conserved r-proteins in various ribosome structures.** Each dot in boxplots represents the measurement of a contacting surface (Å) of either a frequently lost (red boxes) or conserved (blue boxes) ribosomal proteins. Unpaired Wilcoxon test *p*-value was calculated for each structure: PDB ID: 1W2B (*Haloarcula marismortui*, the structure contains 50S subunit and TF), PDB ID: 2ZJR (*Deinococcus radiodurans*, 50S), PDB ID: 4V8X (*Thermus thermophilus*, the complete ribosome), PDB ID: 5DM6 (*Deinococcus radiodurans*, 50S), PDB ID: 5H5U (*Escherichia coli*, the complete ribosome), PDB ID: 6H4N (*Escherichia coli*, the complete ribosome).

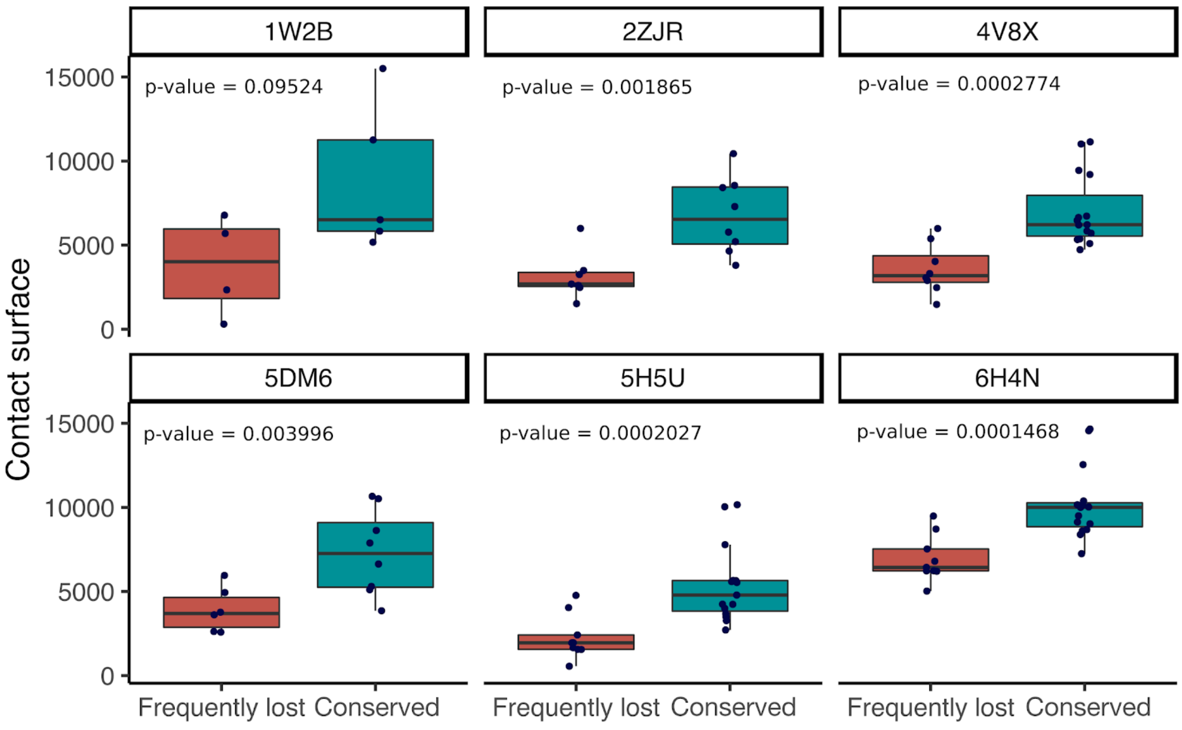

**Figure S5. Deletions in loops of rRNA molecules shown on the ribosome structure (PDB ID: 5H5U, *Escherichia coli*).** Ribosomal proteins are colored in pale yellow, 23S rRNA, in gray, 16S rRNA, in teal. Deleted regions of 23S rRNA are colored in magenta, deletions in 16S rRNA are colored in red. All deletions are labelled according to positions of rRNA molecules from ribosome structure:

Del1 (23S, 153-160 + 166-173)
Del2 (23S, 1715-1727 + 1731-1744)
Del3 (23S, 2207-2210 + 2215-2218)
Del4 (23S, 2789-2805)
Del5 (16S, 76-81)
Del6 (16S, 89-97)
Del7 (16S, 200-207 + 212-216)
Del8 (16S, 837-841 + 845-849)
Del9 (16S, 1127-1135 + 1139-1145)

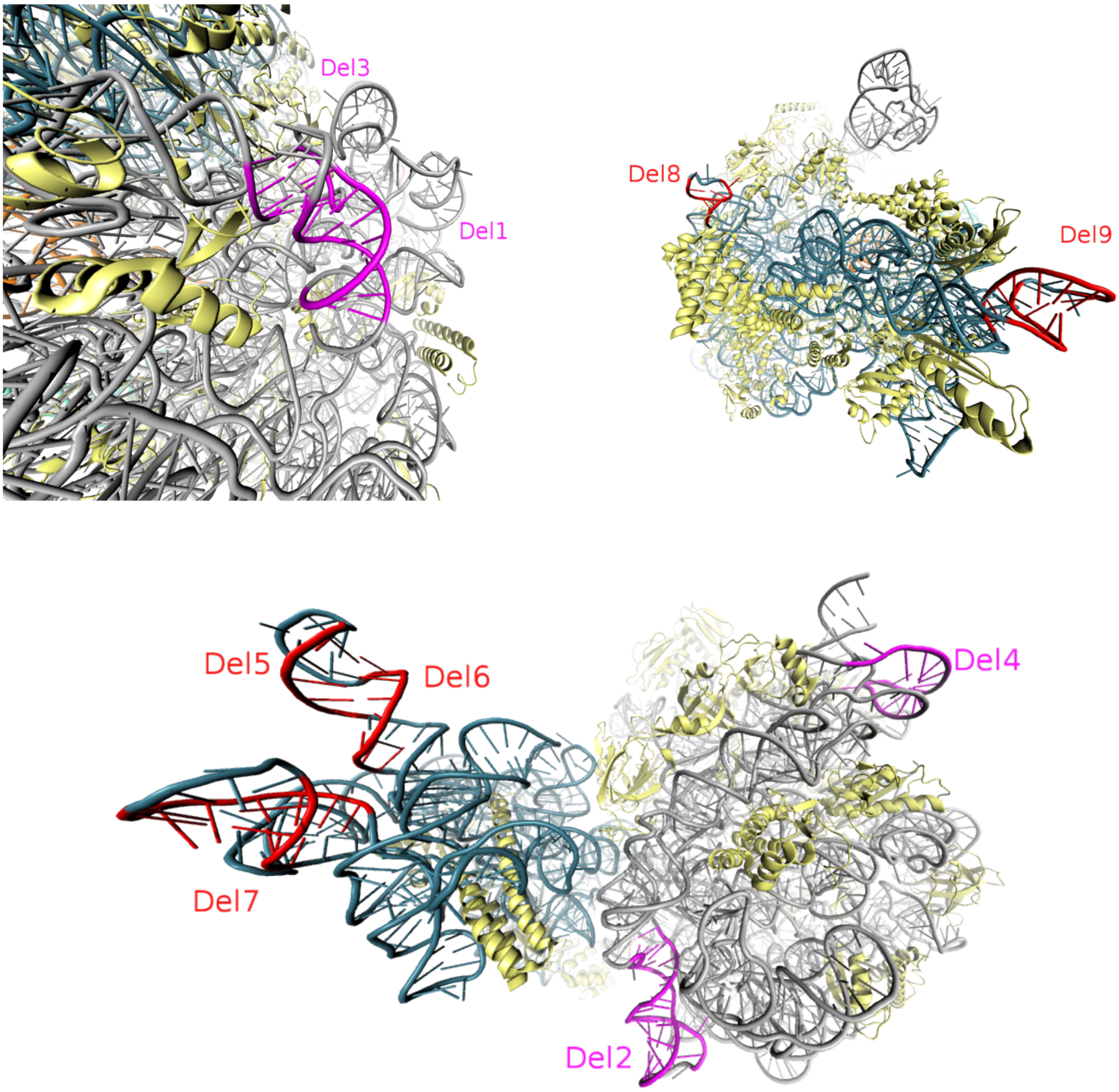

**Figure S6. Loss of r-proteins and the anti-Shine-Dalgarno box.** Dependency between the presence/absence of the anti-SD sequence and the genome length (bp). Each dot represents one strain. The anti-SD sequence is present in 16S rRNA of strains colored blue and absent in strains represented by red dots. The dots are grouped into bars by phyla.

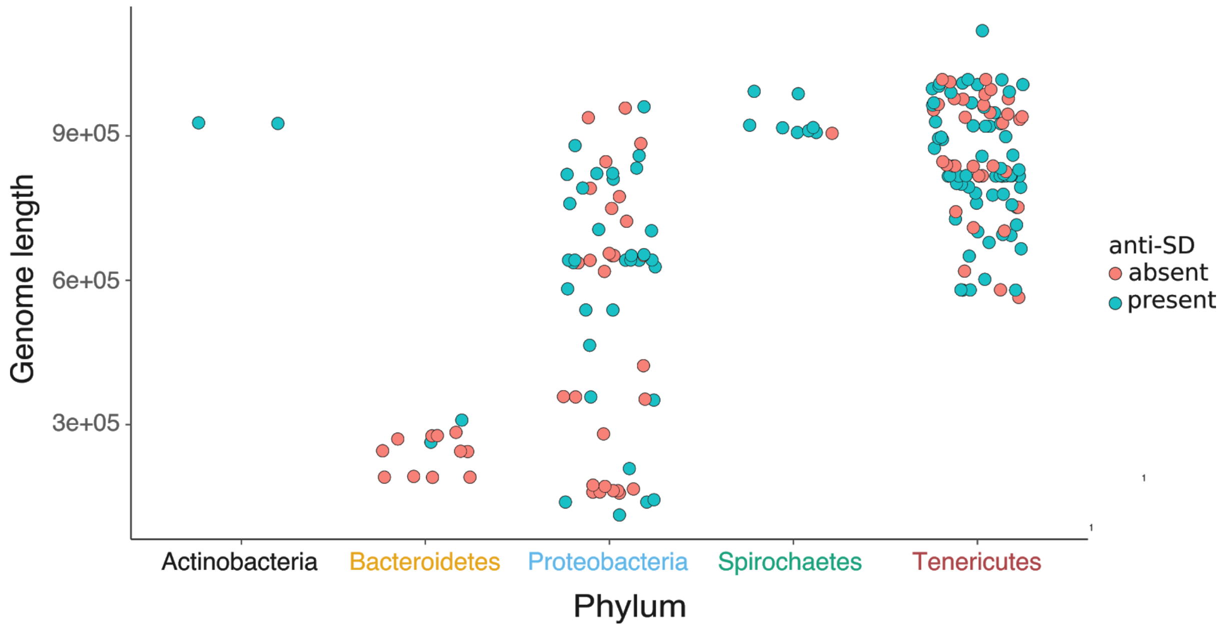

**Table S1.** List of 214 bacterial strains selected for analysis. Strain names of IMG and NCBI databases are given, as well as NCBI Taxonomy and GenBank IDs. See the table in Supplementary Table S1 additional file.

**Table S2.** List of r-proteins and corresponding PFAM domains. R-protein uS3 consists of two domains, however, only the C-terminal domain has been considered as the N-terminal domain is found in many protein families.

| R-protein | Pfam ID | R-protein | Pfam ID |
|---|---|---|---|
| uL1 | PF00687 | bL33 | PF00471 |
| uL2 | PF00181, PF03947 | bL34 | PF00468 |
| uL3 | PF00297 | bL35 | PF01632 |
| uL4 | PF00573 | bL36 | PF00444 |
| uL5 | PF00281, PF00673 | bS1 | PF00575 |
| uL6 | PF00347 | uS2 | PF00318 |
| bL7/L12 | PF00542, PF16320 | uS3* | PF00189, PF07650 |
| bL9 | PF01281, PF03948 | uS4 | PF01479, PF00163 |
| uL10 | PF00466 | uS5 | PF00333, PF03719 |
| uL11 | PF00298, PF03946 | bS6 | PF01250 |
| uL13 | PF00572 | uS7 | PF00177 |
| uL14 | PF00238 | uS8 | PF00410 |
| uL15 | PF00828 | uS9 | PF00380 |
| uL16 | PF00252 | uS10 | PF00338 |
| bL17 | PF01196 | uS11 | PF00411 |
| uL18 | PF00861 | uS12 | PF00164 |
| bL19 | PF01245 | uS13 | PF00416 |
| bL20 | PF00453 | uS14 | PF00253 |
| bL21 | PF00829 | uS15 | PF00312 |
| uL22 | PF00237 | bS16 | PF00886 |
| uL23 | PF00276 | uS17 | PF00366 |
| uL24 | PF17136 | bS18 | PF01084 |
| bL25 | PF01386 | uS19 | PF00203 |
| bL27 | PF01016 | bS20 | PF01649 |
| bL28 | PF00830 | bS21 | PF01165 |
| uL29 | PF00831 | S22 | PF08136 |
| uL30 | PF00327 | TF | PF05697, PF05698 |
| bL31 | PF01197 | bThx | PF17070 |
| bL32 | PF01783 | | |

**Table S3.** Considered parameters of frequently lost and conserved ribosomal proteins: length (in amino acids), evolutionary rate, number of contacts, contacting surface. See the table in Supplementary Table S3 additional file.

**Table S4.** Pagel's test of correlated evolution between the absence of frequently lost proteins and the loss of the anti-Shine-Dalgarno sequence.

| Frequently lost ribosomal protein | Pagel's test of correlated evolution *p-value* (with Bonferroni correction) |
|---|---|
| bL9 | 0.04 |
| uL24 | 1 |
| bL25 | 1 |
| uL29 | 0.23 |
| uL30 | 1 |
| bL32 | 1 |
| bL34 | 1 |
| bL36 | 1 |
| bS21 | 1 |
| TF | $7.55\times10^{-5}$ |

**Table S5.** Alpha (gamma) parameter utilized for phylogenetic tree construction.

| R-protein | Pfam ID | Status | Alpha |
|---|---|---|---|
| uL4 | PF00573 | conserved | 1.058 |
| uL5 | PF00281, PF00673 | conserved | 0.966, 0.966 |
| bL9 | PF01281, PF03948 | lost | 1.332, 1.324 |
| uL13 | PF00572 | conserved | 1.245 |
| uL14 | PF00238 | conserved | 0.940 |
| uL15 | PF00828 | conserved | 0.982 |
| uL16 | PF00252 | conserved | 0.741 |
| bL20 | PF00453 | conserved | 1.047 |
| uL24 | PF17136 | lost | 1.583 |
| bL25 | PF01386 | lost | 1.992 |
| bL27 | PF01016 | conserved | 0.747 |
| uL29 | PF00831 | lost | 1.609 |
| uL30 | PF00327 | lost | 0.817 |
| bL32 | PF01783 | lost | 0.678 |
| bL34 | PF00468 | lost | 0.661 |
| bL36 | PF00444 | lost | 0.794 |
| uS5 | PF00333, PF03719 | conserved | 0.740, 0.740 |
| uS7 | PF00177 | conserved | 0.969 |
| uS8 | PF00410 | conserved | 1.040 |
| uS9 | PF00380 | conserved | 0.678 |
| uS11 | PF00411 | conserved | 0.858 |
| uS13 | PF00416 | conserved | 0.867 |
| uS17 | PF00366 | conserved | 1.004 |
| bS21 | PF01165 | lost | 1.245 |
| TF | PF05697, PF05698 | lost | 3.320, 3.320 |
